# Spontaneous eye behavior is an objective marker of parkinsonian severity

**DOI:** 10.64898/2026.09.24.754183

**Authors:** Ajay K. Verma, Bharadwaj Nandakumar, Quincey V. Feragen, Kit Acedillo, Noah Hjelle, Claudia M. Hendrix, Kelly Duffy, Mark Fiecas, Ying Yu, Biswaranjan Mohanty, Jing Wang, Jerrold L. Vitek, Luke A. Johnson

## Abstract

Eye behavior disruption is reported in Parkinson’s disease (PD), but its correlation with disease severity has not been well examined. Here, in a nonhuman primate model of PD, we show that spontaneous eye behavior is impaired in parkinsonism and correlates with clinical rating scores and subthalamic beta oscillatory activity. These findings suggest that spontaneous eye behavior could serve as an objective, non-invasive biomarker of parkinsonian severity.

---

The severity of Parkinson’s disease (PD), a debilitating neurodegenerative disorder, is typically assessed using a clinician’s rating of the cardinal motor signs—rigidity, bradykinesia, akinesia, tremor, and postural instability^1^. These assessments, however, are subjective, constrained to clinic visits, offer limited temporal resolution, and are prone to inter-rater variability^2^. Excessive beta oscillatory activity (broadly defined 8-35 Hz) in the subthalamic nucleus (STN) is an established biomarker of PD motor symptom severity^3–5^, driving the development of subject-specific, objective beta-based deep brain stimulation (DBS) therapies^3,6,7^. Recording STN beta oscillatory activity requires an invasive DBS electrode implant, limiting its reach to small subsets of patients. Therefore, there is an essential need to develop an objective and non-invasive marker of disease severity applicable to all individuals at risk of PD progression, especially those exhibiting prodromal symptoms of parkinsonism^8^.

The basal ganglia play a central role in motor control^9,10^, and their dysfunction in PD is associated with the manifestation of cardinal motor signs^11,12^. Beyond motor control, the basal ganglia are also involved in regulating eye movements such as saccades and blinks through their anatomical connections to the superior colliculus^13,14^. Consequently, clinical and preclinical studies show that eye behavior is altered in PD^15–20^, and that eye behavior features used in machine learning models can classify parkinsonism and fluctuations in clinical symptoms^21–23^. However, the relationship between eye behavior, pathological beta oscillatory activity in the basal ganglia, and clinical motor scores remains largely unexplored—despite emerging evidence linking beta oscillatory activity to both oculomotor control^24^ and non-motor symptoms^25–28^. Previous studies reporting oculomotor dysfunction in PD employed sophisticated eye-tracking techniques and/or complex behavioral tasks^18,21,29^, limiting their application beyond clinics or laboratory settings.

Advancements in object-tracking techniques have enabled the use of a simple, cost-effective camera to assess eye behavior as a biomarker of parkinsonism^30,31^. Our goal was to investigate whether changes in spontaneous eye behavior, captured using a commercially available off-the-shelf webcam and DeepLabCut^32^, correlate with medication-induced changes in beta oscillatory activity and clinical rating scores. We first characterized in the same subject the effects of parkinsonism on spontaneous eye behavior (saccade and blink rates). Next, we administered the dopaminergic medication levodopa (L-dopa) and recorded spontaneous eye behavior, clinical rating scores, and STN beta oscillatory activity across the time course during which the medication took effect and decayed. We hypothesized that spontaneous eye behavior would be significantly impaired in the parkinsonian state compared to the normal state^18,19^, and that L-dopa would improve spontaneous eye behavior in parallel with the improvements in motor scores and reductions in subthalamic beta power. These insights will establish spontaneous eye behavior as a simple, objective, and non-invasive biomarker for tracking both disease severity and therapy efficacy.

**Figure 1a** illustrates the distributions of saccade rate (saccades/min) and blink rate (blinks/min) in normal and parkinsonian conditions. Moderate parkinsonism reduced saccade and blink rates in all subjects. However, saccade rate, but not blink rate, varied as a function of parkinsonian severity (Subject 1). Moreover, saccade rate demonstrated a larger effect size (Cohen’s d) to distinguish the moderately parkinsonian state from normal in all three subjects **(Figure 1b)**.

Having established that parkinsonism impairs spontaneous eye behavior, we next investigated whether these metrics correlate directly with motor symptoms (mUPDRS) and the power of STN beta oscillatory activity. To test this, we acutely administered L-dopa in parkinsonian NHPs to dynamically modulate motor symptoms and STN beta oscillatory activity. The experimental recording timeline is shown in **Figure 2a**, and an example PSD highlighting alterations in STN low-beta oscillatory activity (8-20 Hz) is shown in **Figure 2b**. The modulation in mUPDRS, saccade rate, blink rate, and STN low-beta power during the time course of medication effect is shown in **Figure 2c**. Next, we tested whether spontaneous eye behavior correlates with mUPDRS and STN beta power. Across subjects, saccade rate correlated negatively with both mUPDRS and STN beta power **(Figure 2d)**. Conversely, the relationships between blink rate, mUPDRS, and STN beta power were not significant across subjects **(Figure 2d)**.

An objective, non-invasive, and cost-effective tool to assess parkinsonian severity remains a key clinical need. This study sought to examine whether spontaneous eye behavior, extracted from webcam footage and using state-of-the-art neural network-based object tracking (i.e., DeepLabCut), can serve as an objective, non-invasive marker of parkinsonian severity. Using a within-subject design in an NHP model of PD, we first show that both saccade rate and blink rate are significantly altered in advanced parkinsonism compared to the normal state (**Figure 1**), aligning with prior studies using sophisticated eye-tracking techniques^14,18,21^.

In both subjects, saccade rate was negatively correlated with mUPDRS and STN beta power, but blink rate correlations with clinical rating scores and STN beta oscillatory activity were not significant across subjects **(Figure 2d)**. These findings suggest that saccade rate, but not blink rate, is a key biomarker of parkinsonian severity. Data from a progressive model of PD (normal, mild, moderate parkinsonian states) further confirmed saccade rate as a key biomarker of parkinsonian severity: saccade rate altered progressively at each severity stage, whereas blink rate only distinguished normal from moderate states (Subject 1, **Figure 1a**). Furthermore, while both saccade rate and blink rate were significantly reduced in a moderately parkinsonian state compared to normal, the saccade rate exhibited a larger effect size than the blink rate, suggesting saccade rate could be a more sensitive marker of parkinsonism than blink rate (**Figure 1b**). Because saccade rate declines early in disease progression and closely tracks severity, it could represent both a sensitive biomarker to guide the timing of disease-modifying interventions and an objective metric for evaluating novel therapeutics in preclinical and clinical settings.

Our finding that saccade rate, but not blink rate, tracks the disease severity suggests that different neural circuits underlie the control of saccades and blinks and warrants further discussion. The basal ganglia output nuclei, especially the substantia nigra pars reticulata (SNr), have strong inhibitory projections to the superior colliculus^33^, which is critically involved in saccadic eye movements^34^. In parkinsonism, the SNr becomes hyperactive, imposing excessive inhibition on the superior colliculus^35–37^, leading to disruption in saccadic eye movements. In contrast, control of blinks relies on a distributed circuit integrating basal ganglia signals with pontine brainstem networks regulating blink circuitry^38^. This suggests that earlier in disease progression, the blinking network may compensate for the loss of dopaminergic signaling and prevent disruption in blink rate, as suggested by our data (**Figure 1a**, Subject 1) and previous studies showing that blink rate may not reflect degeneration of the dopaminergic system^39,40^. While this circuit model remains speculative, simultaneous multisite electrophysiological recordings are warranted to decipher the neural circuits engaged in controlling spontaneous saccades and blinks.

In summary, our data provide evidence for spontaneous eye behavior as a marker of parkinsonian severity, informing the development of a non-invasive, objective, cost-effective, and portable assessment system. Furthermore, these data can serve as an objective marker to assess the efficacy of novel therapies in clinical and preclinical settings for improving parkinsonian symptoms. While limited by the small sample size typical of NHP research, evaluating eye movement dynamics across disease progression provides unique preclinical insight rarely feasible in human studies. Although we characterized dysfunction in eye movements, future electrophysiological studies are essential to improve mechanistic understanding of the disruption in eye behavior that is associated with progression of parkinsonism.

**Figure 1.**
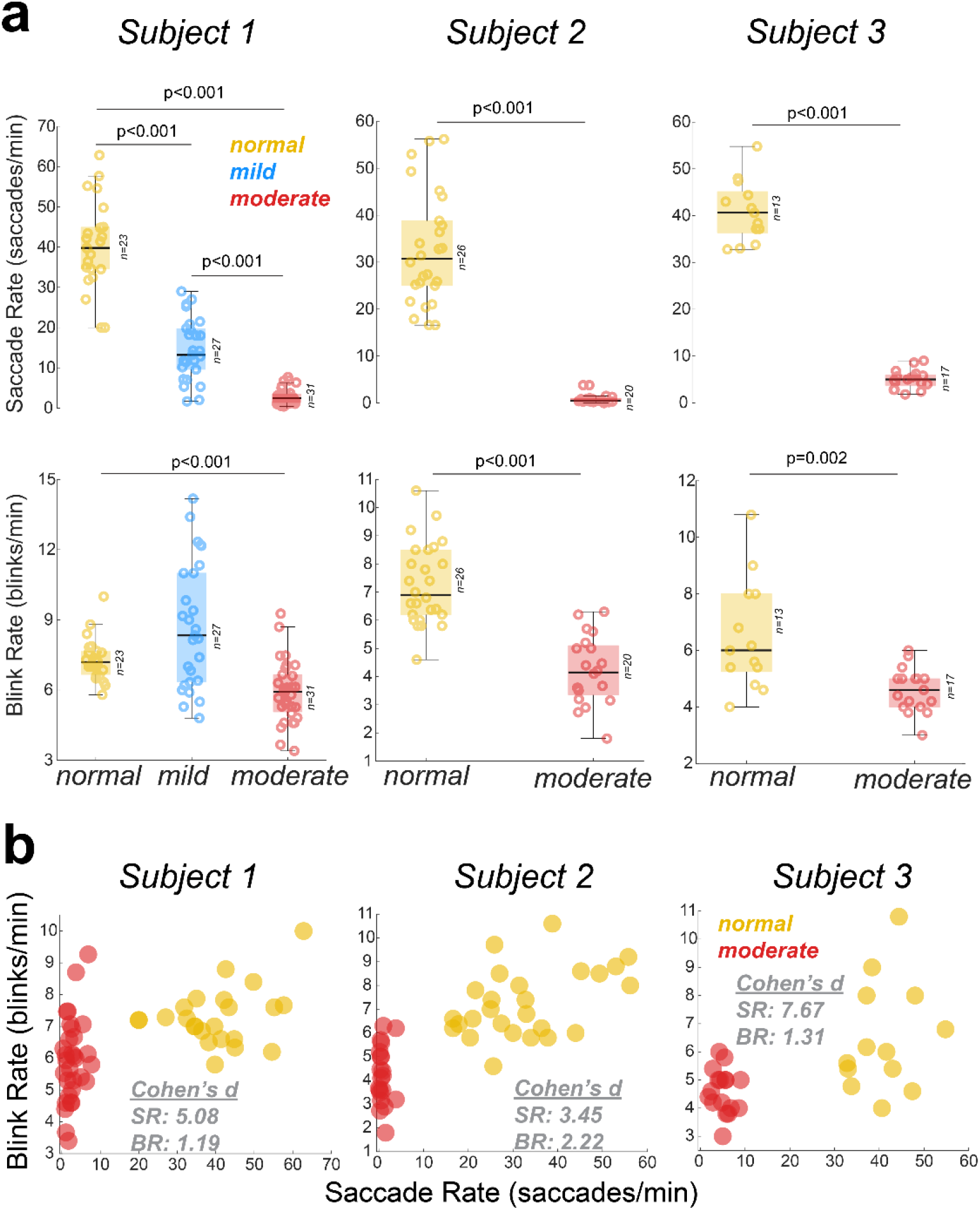
The saccade rate (SR) and blink rate (BR) were significantly altered in the moderately parkinsonian state compared to the normal state in all 3 subjects **(a).** Saccade rate, but not blink rate, tracked progression of parkinsonsim (Subject 1). The saccade rate and blink rate scatter plot exhibit the potential of eye behavior features for differentiating normal and parkinsonian states **(b)**. Based on effect size, saccade rate is a more robust marker of parkinsonism compared to blink rate **(b)**.

**Figure 2.**
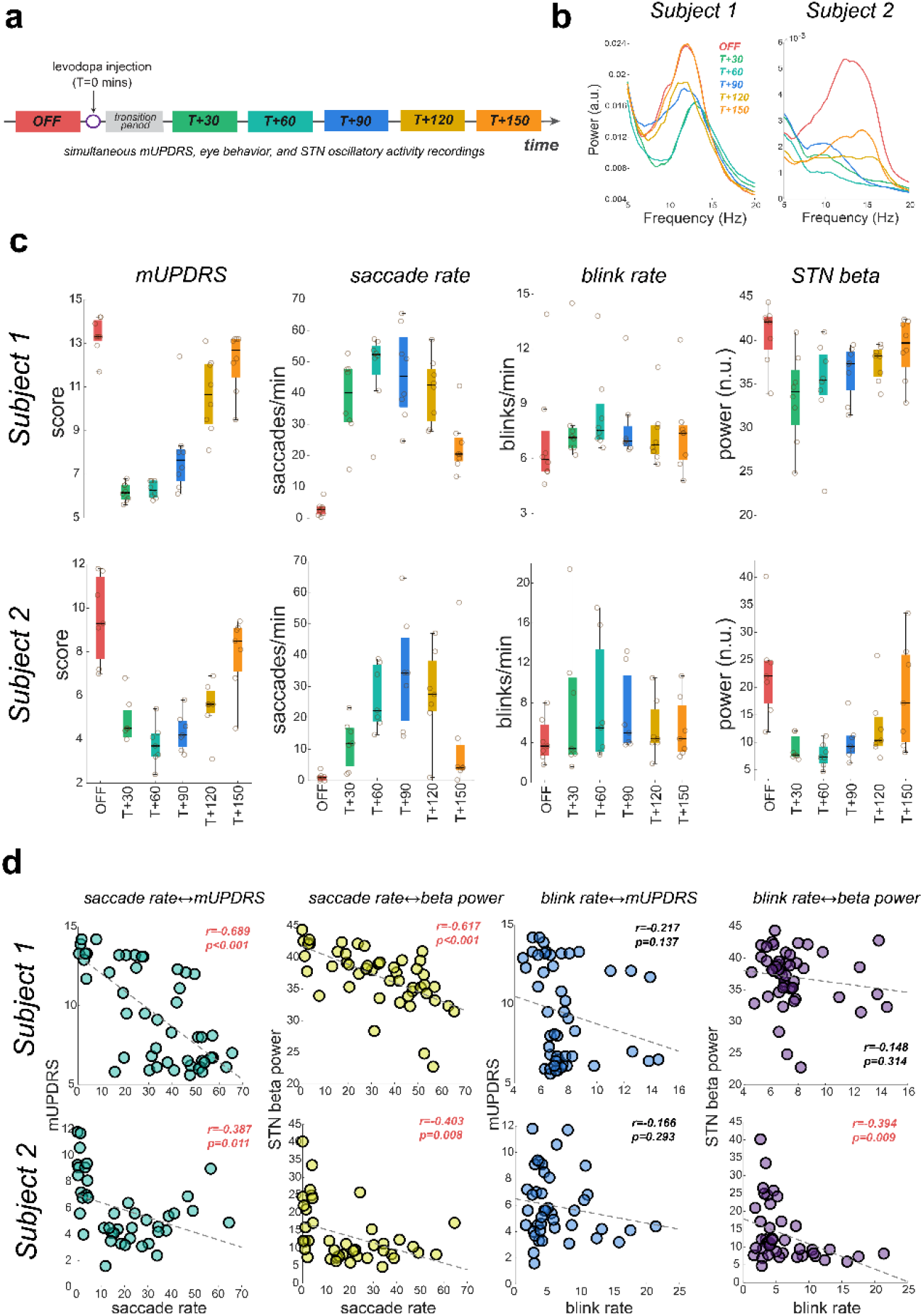
The experimental timeline for L-dopa time course data recordings is shown in **a.** An example PSD highlighting changes in STN beta oscillatory activity is shown in **b**. The distributions of mUPDRS, saccade rate, blink rate, and STN beta power are shown in **c**, while the correlations between eye-behavior features and subjective (mUPDRS) and objective (beta power) measures of parkinsonian severity are shown in **d**. Saccade rate correlated with mUPDRS and beta power in both subjects, but the correlation of blink rate with mUPDRS and beta power was not significant across subjects **(d)**.

## Methods

### Experimental protocol and data collection

All procedures were approved by the University of Minnesota Institutional Animal Care and Use Committee and complied with the US Public Health Service policy on the humane care and use of laboratory animals. Three aged female rhesus macaques NHPs (Subject 1: 27 years; Subject 2: 22 years; Subject 3: 16 years) were used in this study.

#### Experiment 1

The goal was to examine whether the parkinsonian state alters spontaneous eye behavior. To investigate this, spontaneous eye movements were recorded using a USB video camera (20 frames/s) to extract and compare saccade and blink rates between normal and parkinsonian states.

The recordings were first performed in the normal state, followed by recordings in the parkinsonian state. The subjects were rendered parkinsonian through intracarotid or intramuscular injections (0.2–0.4 mg/kg) of the neurotoxin 1-methyl-4-phenyl-1,2,3,6-tetrahydropyridine (MPTP). The MPTP was administered following protocols established by our research center, adapted from Masilamoni et al^41^. The severity of parkinsonian motor signs was assessed using a modified Unified Parkinson’s Disease Rating Scale (mUPDRS)^42^. This scale rates axial motor symptoms (gait, posture, balance, defense reaction) and limb symptoms (rigidity, bradykinesia, akinesia, tremor) from 0 (normal) to 3 (severe), yielding a maximum total score of 39^43^. Based on these scores, parkinsonian severity was defined as mild (3-15), moderate (18-31), or severe (29-39). The mUPDRS scores were recorded continuously to ensure the parkinsonian motor signs remained stable. This experiment reports data from normal, mild, and moderately parkinsonian conditions in one subject and normal and moderately parkinsonian conditions in two subjects.

#### Experiment 2

This experiment investigated whether spontaneous eye behavior correlates with STN beta oscillatory activity and clinical rating scores. To modulate beta power and mUPDRS scores, the dopaminergic medication levodopa (L-dopa) was administered in two subjects (Subject 1: 10 mg/kg; Subject 2: 15 mg/kg). Throughout the onset and decay of the medication’s effects, spontaneous eye behavior, mUPDRS scores, and STN oscillatory activity were recorded via implanted DBS electrodes^25^. The specific L-dopa doses were determined based on prior characterizations of the subjects’ motor responses, selecting the lowest dose that improved parkinsonian motor signs without inducing side effects (e.g., dyskinesia or nystagmus). The data collection experimental protocol is shown in **Figure 2a**. Due to experimental constraints, axial motor signs were not assessed in Experiment 2, resulting in a maximum mUPDRS score of 27, with mUPDRS scores 3-9, 10-18, and 19-27 representing mild, moderate, and severe parkinsonian states, respectively. Experiment 2 was conducted only in two subjects (Subjects 1 and 2).

### Data Analysis

#### DeepLabCut (DLC) training and eye behavior extraction

Eye behavior was tracked using a 20 frames/s video camera. To extract pupil and eyelid movements, a ResNet-50 convolutional neural network was trained using DeepLabCut^32^. The training set consisted of 20 videos averaging 180,000 frames. From this set, a 300-frame subset was manually labeled to annotate the right pupil, left pupil, upper right eyelid margin, and lower eyelid margin. The labeled dataset was shuffled, split into training and testing sets (95:5 ratio), and trained for 10,000 iterations. After evaluating the network on an independent test dataset, the extracted markers were visually confirmed, and any tracking outliers were manually corrected.

The DeepLabCut output provides the horizontal (x) and vertical (y) coordinates for the pupils, lower eyelid margins, and upper eyelid margins within the field of view (FOV). These position vectors were re-centered for every video by aligning the pupil position with the center of the FOV. Next, the velocity of the pupil and eyelid movements was quantified using a discrete central difference approximation of the derivative.

Saccades were identified when the resultant pupil velocity exceeded a threshold of 5 times the median absolute deviation of the resultant velocity across the entire recording^44^. If position traces contained noise from animal jitter, saccade identification was visually verified, and the threshold was increased to prevent false positives. Blinks were identified when eyelid movement velocity exceeded a threshold of 0.2, provided the blink duration lasted between 0.1 and 0.5 seconds. Blinks longer than 0.5 seconds were classified as transitions between wake and sleep states. Finally, saccades were extracted exclusively during non-blink, awake epochs (i.e., eyes open).

### Local field potential (LFP) analysis and spectral power extraction

Subject 1 was instrumented with an 8-ring DBS lead (0.5 mm contact height, 0.5 mm inter-contact spacing, 0.625 mm diameter; NuMed Inc.) while subject 2 was instrumented with a segmented DBS lead (4 rings, 3 contacts per ring; 0.5 mm contact height, 0.5 mm spacing between rows, 0.8 mm diameter; Heraeus). The contact leads targeting the STN were chosen based on post-surgical registration of CT images with pre-surgery MRI. STN LFPs were sampled using a TDT neural data acquisition system (Tucker Davis Technologies) at 24000 Hz.

STN LFPs for subject 1 were derived as the differential signal between vertically adjacent contacts on the implanted DBS leads; for subject 2, the three contacts per ring were averaged prior to signal differentiation between vertically adjacent rings. For spectral analysis, the acquired data were downsampled to 200 Hz. The trial-averaged power spectral density (PSD) was then computed using the Chronux toolbox in MATLAB^45^, selected for its robust anti-leakage characteristics and optimal time-frequency resolution trade-off. Specifically, a trial-averaged power spectral density (2-second epochs, 5 Slepian multitapers, time-bandwidth product of 3) was calculated for all epochs when the animal was awake (i.e., eyes open). The spectral power in the low-beta band was obtained by integrating the PSD across the 8–20 Hz band and subsequently normalized to the total power (1–100 Hz).

#### Statistical Analysis

All statistical tests were performed independently for each subject; data were not pooled across subjects. The Wilcoxon rank-sum test was used to evaluate changes in eye behavior features between normal and parkinsonian states, and Cohen’s d was calculated to determine effect size. To assess how eye behavior changed as parkinsonism progressed (i.e., normal, mild, moderate), the Kruskal-Wallis test was applied, followed by Tukey’s post hoc correction. The Pearson correlation coefficient was used to evaluate the degree of correlation between variables. All test results were considered statistically significant at p<0.05. All statistical tests were performed in MATLAB 2024b.

## Authors contribution

A.K.V., B.N., and L.A.J. conceived this study. A.K.V., B.N., and L.A.J. designed the experiment. A.K.V., B.N., and Q.V.F. performed the literature review and wrote the first draft. B.N. analyzed data. A.K.V. made figures. A.K.V., K.D., and M.F. performed statistical analysis. K.A., N.H., A.K.V., B.N., Y.Y., and B.M. collected data. A.K.V., B.N., Q.V.F., C.M.H., J.W., J.L.V., and L.A.J. interpreted data. Y.Y., N.H., and B.M. performed animal care. All authors critically reviewed the manuscript and approved it for publication.

## Conflict of interest

JLV serves as a consultant for Medtronic, Boston Scientific, and Abbott. He also serves on the Executive Advisory Board for Abbott and is a member of the scientific advisory board for Surgical Information Sciences. JLV has no non-financial conflicts to disclose. The remaining authors have no competing interests to disclose.

## Acknowledgment

This work was supported by the National Institutes of Health, National Institute of Neurological Disorders and Stroke (NINDS): R01-NS110613, R01-NS131371, R01-NS058945, R01-NS037019, R37-NS077657, P50-NS123109. This research was supported by the National Institutes of Health’s National Center for Advancing Translational Sciences (NCATS), grants K12TR004373 and 1UM1TR004405. The content is solely the responsibility of the authors and does not necessarily represent the official views of NINDS or NCATS. MnDRIVE (Minnesota’s Discovery, Research, and Innovation Economy) Brain Conditions Program, MNREACH, and philanthropic support from the Engdahl Family Foundation.

## Data availability

The data supporting the findings of this study are available from the corresponding author upon reasonable request.

